# PTBP2-dependent alternative splicing regulates protein transport and mitochondria morphology in post-meiotic germ cells

**DOI:** 10.1101/497461

**Authors:** Molly M. Hannigan, Hisashi Fujioka, Adina Brett-Morris, Jason A. Mears, Donny D. Licatalosi

## Abstract

The RNA binding protein PTBP2 (polypyrimidine tract binding protein 2) is a key regulator of tissue-specific alternative RNA splicing. In the testis, PTBP2 is expressed in meiotic and post-meiotic germ cells (spermatocytes and spermatids, respectively). In these cells, PTBP2 is required for proper alternative mRNA splicing for over 200 genes, disproportionately affecting genes encoding proteins involved in protein trafficking via transport vesicles. In this study, we used electron microscopy to test the hypothesis that protein trafficking is impaired in the absence of PTBP2, and to further investigate why spermatogenesis abruptly ceases in PTBP2-deficienct spermatids. Ultrastructural analysis shows that protein trafficking in spermatids is aberrant in the absence of PTBP2. Unexpectedly, we also found that mitochondria morphology and number are significantly altered in PTBP2-deficient spermatids, consistent with increased mitochondria fission. Furthermore, we show that genes with key roles in mitochondria dynamics and function are post-transcriptionally regulated by PTBP2 and in different stages of spermatogenesis. Collectively, the data provide ultrastructural evidence that alternative splicing regulation by PTBP2 during spermatogenesis is critical for proper regulation of protein trafficking and mitochondria morphology.

## INTRODUCTION

Alternative exon splicing (AS) allows individual genes to produce multiple mRNA isoforms with overlapping but distinct protein-coding and/or non-coding sequences. As a result, AS mRNA isoforms can encode proteins with distinct biochemical properties, or have different patterns of mRNA localization, stability, or translation. AS isoforms are highly regulated across different tissues and stages of development. Interestingly, exons spliced in a tissue-restricted manner frequently encode intrinsically disordered regions (IDRs), linear peptide motifs, and post-translational modification sites. These have been proposed to tune protein function, and rewire protein-protein interaction networks and signaling pathways (Merkin et al., 2012; Buljan et al., 2012; Ellis et al., 2012; Keul et al., 2018). While it is known that nearly all multi-exon protein coding genes in mice and humans generate multiple AS mRNA isoforms (Barbosa-Morais et al., 2012; Merkin et al., 2012), our understanding of how networks of AS isoforms are coordinately regulated to alter cellular properties remains limited.

The testis expresses considerably more AS isoforms compared to other tissue types (Clark et al., 2007; Yeo et al., 2004; Soumillon et al., 2013). Within the testis are seminiferous tubules, which is where spermatogenesis occurs. Throughout their development, germ cells are in direct contact with Sertoli cells that provide nutritional and structural support to spermatogenic cells as they translocate from the basal membrane to the lumen of the seminiferous epithelium (Griswold, 1998). The first phase of spermatogenesis involves a series of cell proliferation events in which spermatogonia can self-renew to maintain a stem cell pool, or proliferate and differentiate to yield a large pool of cells that enter meiosis. In meiotic cells (spermatocytes), DNA replication is followed by homologous recombination between non-sister chromatids, and two successive cell divisions to produce haploid round spermatids. In the last phase, post-meiotic spermatids undergo a differentiation process called spermiogenesis before they are released into the lumen as spermatozoa. This process involves dramatic morphological changes including compaction of the nucleus, development of a flagellum, and formation of the acrosome – an exocytotic vesicle that contains hydrolytic enzymes required for oocyte fertilization.

Strikingly, two spermatogenic cell types (spermatocytes and spermatids) are responsible for most of the AS isoform complexity found in the testis (Soumillon et al., 2013). The functions and regulation of AS in these and other spermatogenic cells is just beginning to be understood (O’Bryan et al., 2013; Schmid et al., 2013; Iwamori et al., 2016; Hannigan et al., 2017). Using available RNA-sequencing (RNA-seq) data from spermatogonia, spermatocytes, round spermatids, and spermatozoa (Soumillon et al., 2013), we previously identified 1030 stage-specific changes in AS mRNA isoforms as germ cells move through the spermatogenic pathway (affecting expression of 912 genes) (Hannigan et al., 2017). The majority of these AS differences (75%) exist between mitotic and meiotic cells (spermatogonia and spermatocytes, respectively). Few AS changes occur as spermatocytes transit through meiosis to produce round spermatids, and these were generally small in magnitude. In contrast, greater differences in AS isoforms exist between mRNAs present in round spermatids compared to spermatozoa. These patterns indicate that AS is subject to distinct stage-specific programs of regulation during spermatogenesis.

We previously showed that the tissue-restricted AS regulator PTBP2 (polypyrimidine tract binding protein 2) is required for spermatogenesis (Zagore et al., 2015). PTBP2 is a member of the PTB family of RNA binding proteins (RBPs) and has key roles in developmental regulation of AS in the nervous system (Licatalosi et al., 2012; Zheng et al., 2012; Li et al., 2014). In the testis, PTBP2 expression is restricted to germ cells, with the highest levels in spermatocytes and round spermatids (Xu and Hecht, 2007). Ablation of *Ptbp2* expression using a conditional gene knockout approach (cKO) resulted in spermatogenic failure with apoptosis of a subset of spermatocytes and complete arrest of spermatids as giant multinucleated cells during the first few steps of differentiation (Zagore et al., 2015). RNA-Seq analysis of cKO and wild type (WT) testes indicated that PTBP2 is required for proper AS regulation of 217 genes, and has important roles in stage-specific AS regulation during spermatogenesis (Hannigan et al., 2017). Integrating the RNA-Seq data with transcriptome-wide mapping of *in vivo* PTBP2-RNA interactions demonstrated that PTBP2 primarily functions in germ cells to directly repress splicing of select AS exons in via RNA-binding near the 3’ splice site, as in the brain (Hannigan et al., 2017; Licatalosi et al., 2012). Thus, PTBP2 is a key regulator of AS isoform expression in spermatogenesis.

Our understanding of why loss of PTBP2-dependent AS regulation leads to spermatogenic failure is limited. Examination of the affected genes showed a strong enrichment for those whose protein products have roles in GTPase-mediated signaling and protein sorting via transport vesicles, suggesting that these processes may be perturbed in *Ptbp2* cKO germ cells. Indeed, testes with *Ptbp2* cKO germ cells had excessive tracks of filamentous actin in neighboring Sertoli cells (Hannigan et al., 2017), an indicator of defective germ cell polarity (Tanwar et al., 2012). Notably, disorganization of the Sertoli cell cytoskeleton was observed prior to the formation of multinucleated spermatids. Altogether, these observations suggest that PTBP2-dependent AS is required to ensure proper protein trafficking within germ cells. They also indicate that imbalances in AS isoforms in germ cells alters cytoskeletal organization in Sertoli cells and their ability to support post-meiotic spermatids.

In this study, we sought to further understand the biological function of PTBP2 in spermatogenesis. To do so, we performed an ultrastructural analysis of WT and *Ptbp2* cKO testes using transmission electron microscopy (TEM), with an emphasis on round spermatids (the most advanced germ cell stage present in *Ptbp2* cKO mice). In particular, we focused on acrosome formation, which is heavily dependent on the trafficking and fusion of vesicles derived from the Golgi apparatus and the endosomal-sorting pathway (Berruti and Paiardi, 2015; Berruti, 2016). Indeed, our TEM analysis indicates these processes are perturbed in the absence of PTBP2. Notably however, defects in acrosome biogenesis in *Ptbp2* cKO mice are distinct from the acrosomal defects observed in other gene knockout mice. Unexpectedly, TEM also revealed abnormally shaped mitochondria in *Ptbp2* cKO spermatids, consistent with increased fission. Indeed, we found that DRP1, a major regulator of mitochondrial dynamics, was significantly elevated in *Ptbp2* cKO testes despite the absence of differences in mRNA encoding DRP1 between WT and cKO testes. We also identified a large set of genes whose AS mRNA isoforms are controlled in a PTBP2-dependent and/or stage-specific manner and encode proteins with roles in regulating mitochondrial dynamics and/or activity. Collectively, our analyses provide important insights into the functions of PTBP2 and AS in regulating protein trafficking and mitochondrial dynamics in post-meiotic germ cells.

## RESULTS

### Acrosome formation is impaired in PTBP2-deficient spermatids

Proper acrosome biogenesis is a key event during the differentiation of round spermatids into spermatozoa and is essential for fertility (Kierszenbaum et al., 2007; Berruti, 2016). Acrosome formation depends on continuous trafficking and fusion of coated vesicles derived from the Golgi apparatus and the endosomal-sorting pathway. During the Golgi phase of acrosome biogenesis, numerous proacrosomal granules derived from trans-Golgi stacks accumulate and fuse to form a large acrosomal granule that migrates towards the nucleus. This granule becomes anchored to the nucleus via the acroplaxome (a keratin and actin-rich scaffold) (Kierszenbaum and Tres, 2004). During the cap phase, the acrosomal granule dramatically increases in size through continuous of addition of Golgi-derived and endosomal-sorted cargo as it flattens and spreads over the nucleus (Berruti, 2016). In WT testes collected at P25 or P30, both Golgi and cap phase spermatids were abundant and readily observed (Figure 1A, B). Consistent with the ages examined, more advanced spermatids were rare or absent (acrosome and maturation phases, respectively). In contrast, acrosomes in the majority of round or multinucleated spermatids had not advanced beyond the Golgi phase in *Ptbp2* cKO testes (Figure 1C). Additional abnormalities in acrosome biogenesis were also observed by TEM. Within multinucleated cells, we frequently observed a single acrosomal granule attached to the nuclear membrane of multiple nuclei (Figure 1D). Acrosome formation defects were also observed in round spermatids that were not multinucleate. Figure 1E shows a representative example with defects consistent with aberrant vesicle fusion (Heimann et al., 1991; Gioria et al., 2017), evidenced by two acrosomal granules in contact with the nuclear membrane and the accumulation of numerous proacrosomal granules nearby. Strikingly, one of the two acrosomal granules in contact with the nuclear membrane appears to have a ruptured membrane, with leakage of acrosomal contents into the cytosol (Figure 1F).

**Figure 1.**
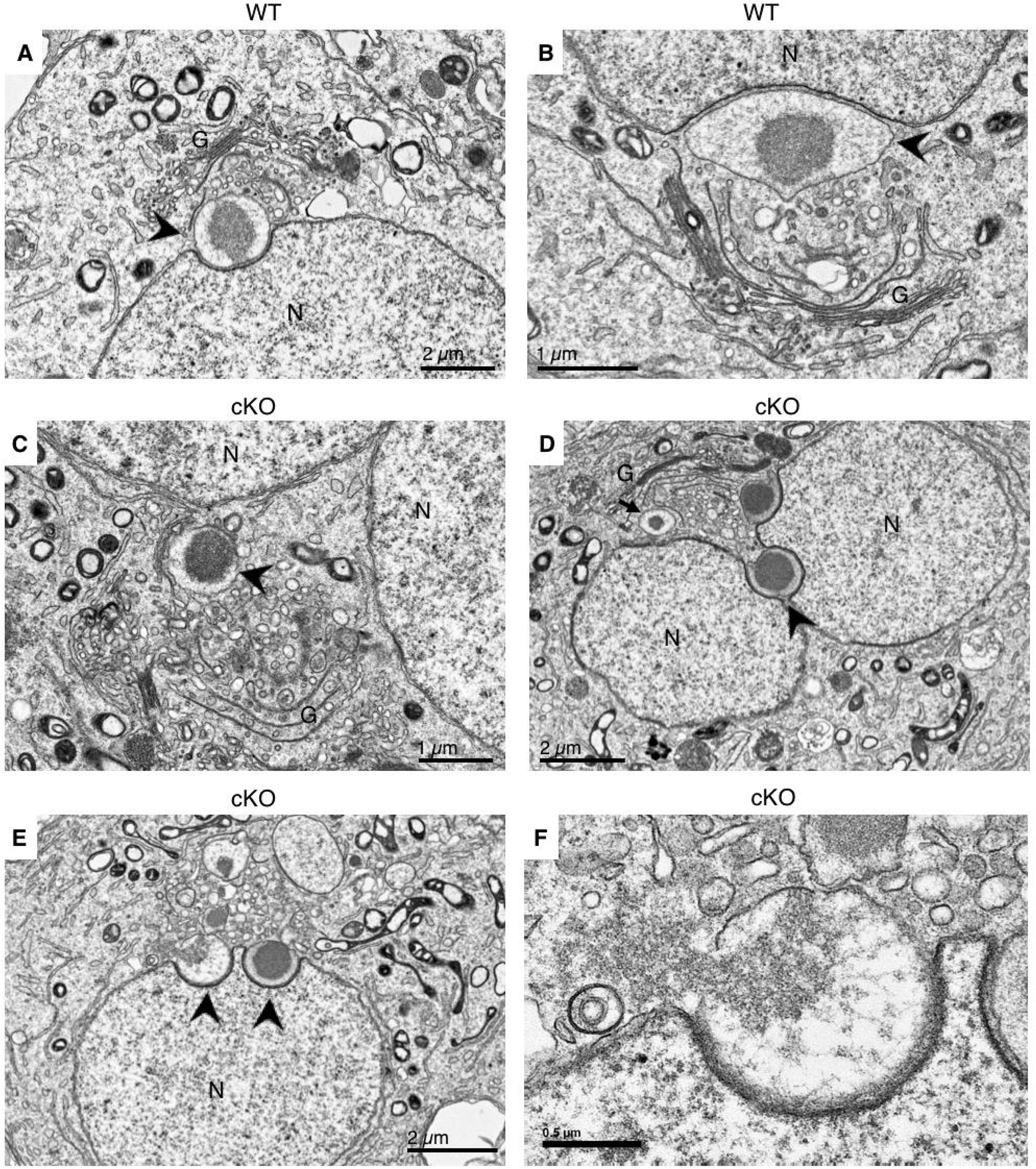
Most round and multinucleate *Ptbp2* cKO spermatids contain only Golgi phase acrosomes. Electron micrographs of WT (A, B) and *Ptbp2* cKO (C-F) spermatids. In WT mice, spermatids in the Golgi (A) and cap (B) phases are readily observed. C) A single proacrosomal granule in close proximity to two nuclei of a multinucleated cKO spermatid. D) A multinucleated cKO spermatid, showing a single acrosome (arrowhead) attached to two nuclei. A large proacrosomal granule is also present (arrow). E) A cKO spermatid with two acrosomes attached to a single nucleus, and several proacrosomal granules in the vicinity. F) Higher magnification of the left acrosome from panel E. Note the discontinuity of the acrosomal outer membrane, and continuity of the acrosomal matrix between the acrosome and cytoplasm. Black arrowhead, acrosome; N, nucleus; G, Golgi apparatus.

In *Ptbp2* cKO testes at P25 and P30, we found few instances in which acrosome formation progressed to the cap phase, when the acrosome increases in size and begins to flatten along the nuclear membrane. Moreover, the few cap phase *Ptbp2* cKO spermatids identified had abnormal features. In one example, a *Ptbp2* cKO round spermatid had multiple coated vesicles within the acrosome (Figure 2A, B). In another, a large membrane-bound vesicle containing smaller vesicles was present within the acrosome (Figure 2C, D). Although coated vesicles are known to participate in acrosome formation, these vesicles immediately shed their clathrin coats after fusion (Bonifacino and Glick, 2004; Huang and Ho, 2006). The accumulation of coated vesicles within the acrosome of cKO spermatids is consistent with aberrant protein transport via vesicle trafficking in cKO spermatids due to loss of PTBP2-dependent AS regulation.

**Figure 2.**
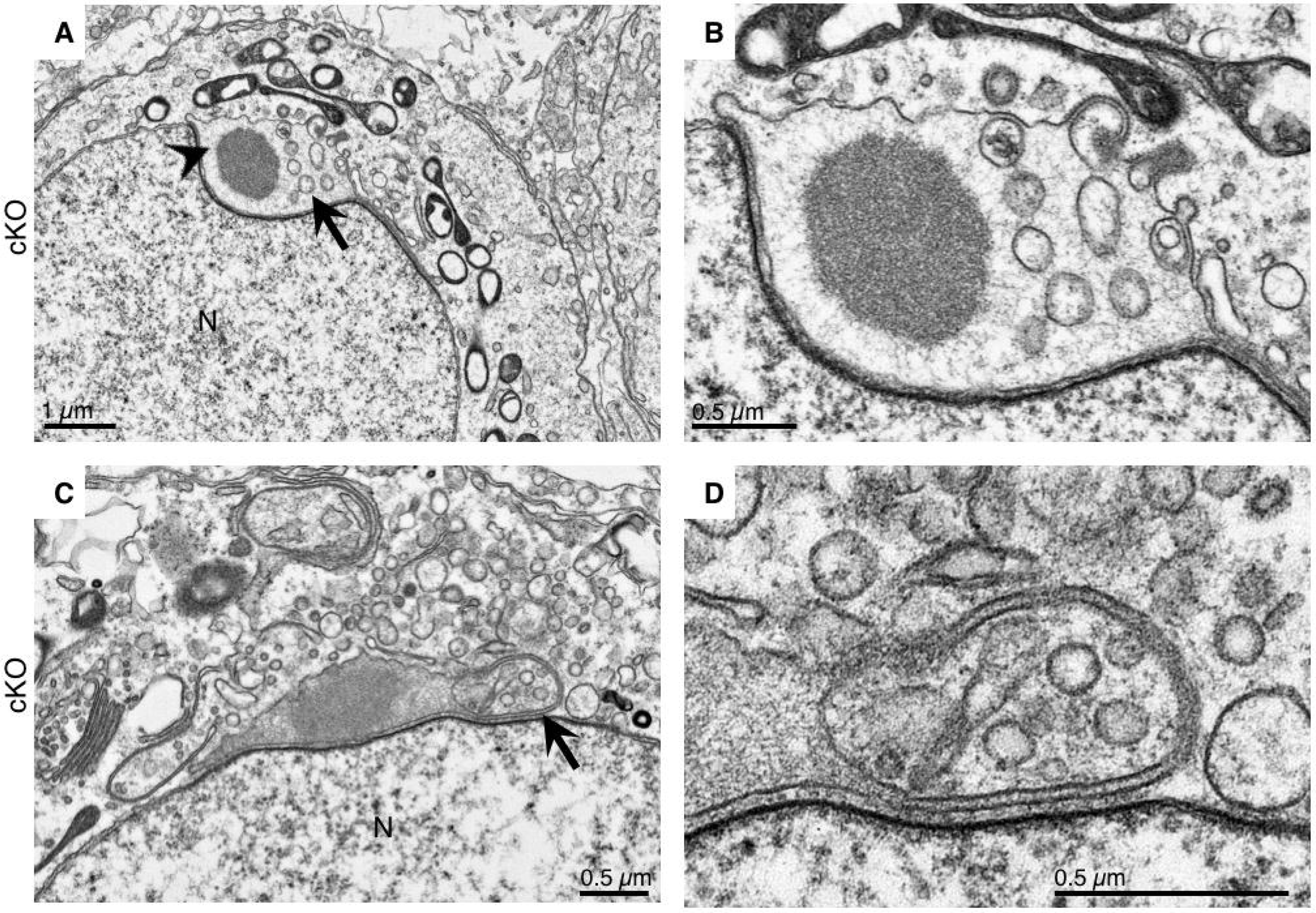
Coated vesicles accumulate within cap phase acrosomes in *Ptbp2* cKO spermatids. EM micrographs of *Ptbp2* cKO spermatids with multiple coated transport vesicles within the acrosome. Panels B and D are higher magnification views of the spermatids shown in panels A and C, respectively.

Altogether, TEM analysis indicates that protein trafficking via transport vesicles is altered in *Ptbp2* cKO spermatids, consistent with our previous observation that PTBP2-dependent AS disproportionately impacts genes involved in protein transport during spermatogenesis.

### PTBP2-deletion alters mitochondrial dynamics in spermatids

During our TEM examination of acrosome formation, we observed additional ultrastructural differences between WT and *Ptbp2* cKO spermatids. The most consistent and prominent of these differences was the number, distribution, and morphology of mitochondria. During spermatogenesis, mitochondria change in morphology and localization dependent on the spermatogenic cell type (Meinhardt et al., 1999). Traditionally, spermatogonia contain orthodox, ovoid-shaped mitochondria with lamellar cristae and electron translucent matrix. During the early stages of meiosis, mitochondria elongate and increase in number, congregating near the outer nuclear membrane. Mitochondria return to a small, round shaped organelle without a clustered arrangement in mid-pachytene spermatocytes through early post-meiotic spermatids. Whereas WT spermatids contained mitochondria with the expected morphology (Figure 3A-C), mitochondria were elongated and dumbbell shaped in *Ptbp2* cKO spermatids (Figure 3D-F). Furthermore, these abnormally shaped mitochondria were much more abundant and clustered in cKO spermatids, compared to the less numerous and more evenly distributed mitochondria seen in WT spermatids. Interestingly, no obvious differences were observed between mitochondria in WT and *Ptbp2* cKO spermatocytes (data not shown).

**Figure 3.**
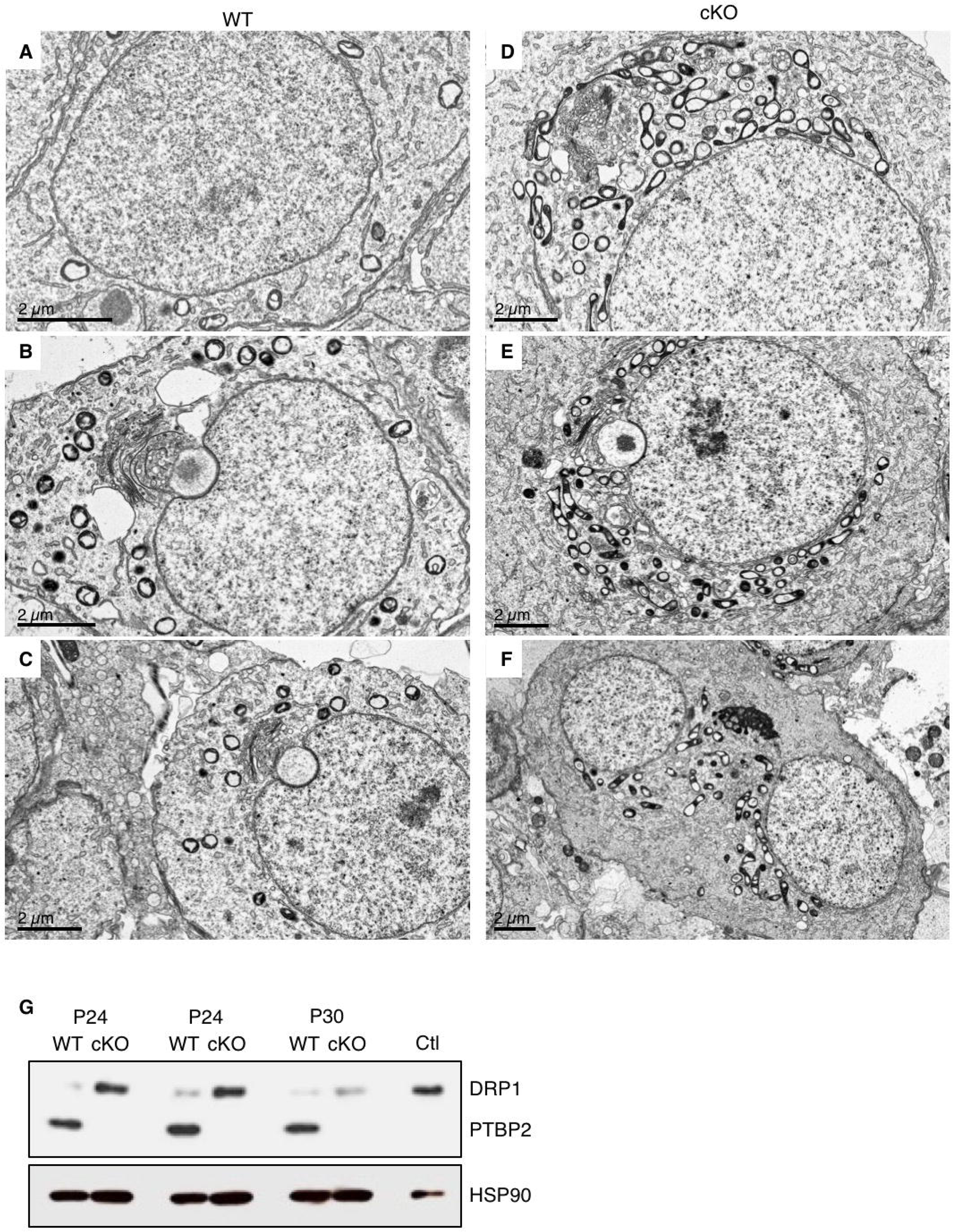
Increased mitochondrial fission in spermatids lacking PTBP2. A-C) EM micrographs of WT spermatids with spherical and evenly distributed mitochondria. D-F) EM micrographs of *Ptbp2* cKO spermatids, which have an accumulation of elongated, dumbbell shaped mitochondria, suggesting these organelles are undergoing fission. G) Western blot analysis showing increased levels of DRP1 in testis lysates from *Ptbp2* cKO mice collected at P24 and P30 compared to WT littermates. Levels of PTBP2 and HSP90 (loading control) are also shown. Ctl corresponds to lysates from HeLa cells with over-expression of DRP1.

Mitochondria morphology, distribution and function are dependent on the balanced regulation of mitochondrial fusion and fission (El-Hattab et al., 2018). The striking increase in the amount of mitochondria and their elongated appearance, suggest that excessive mitochondrial fission may occur in *Ptbp2* cKO spermatids. A main regulator of mitochondrial fission is Dynamin-related protein 1 (DRP1), a critical GTPase that is recruited to the mitochondria and oligomerizes to drive constriction during mitochondrial division (Tilokani et al., 2018). Western blot analysis showed that DRP1 was significantly increased in *Ptbp2* cKO versus WT testis (Figure 3G). Importantly, *Dnm1l* (the gene encoding DRP1) and other critical mediators of mitochondrial dynamics had similar RNA levels in WT and cKO testes (Table 1) suggesting post-transcriptional up-regulation of factors involved in mitochondrial metabolism in the absence of PTBP2. Taken together, these observations suggest that increased DRP1 may increase mitochondrial fission in cKO spermatids, resulting in differences in the number and morphology of mitochondria between WT and *Ptbp2* cKO spermatids.

**Table 1.**
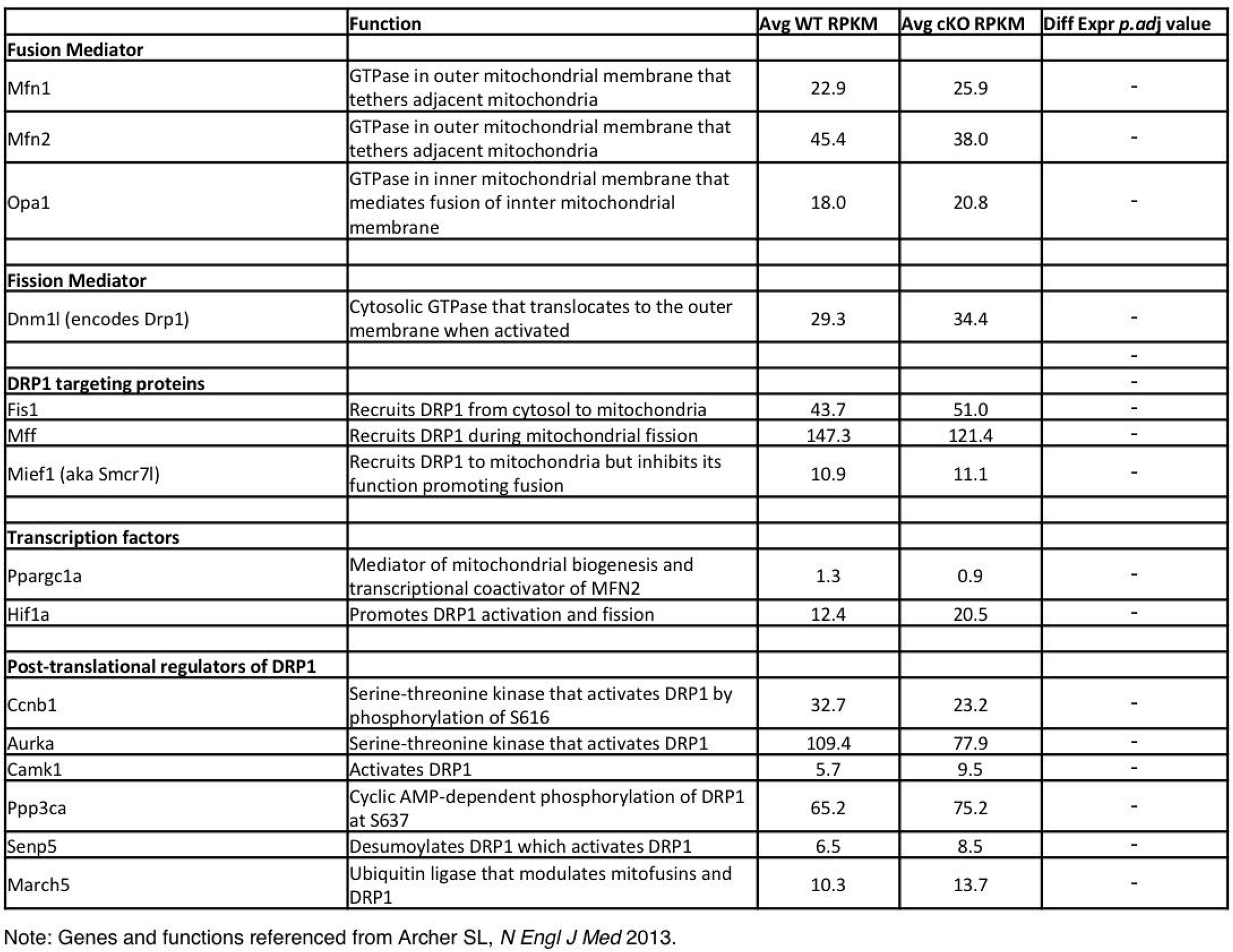
Gene expression of factors involved in mitochondria dynamics. List of factors that function in mitochondrial dynamics and their corresponding gene expression values in WT and *Ptbp2* cKO testes. As displayed by the right most column, none of these factors have differential gene expression in the absence of PTBP2. These genes and functions are referenced from (Archer, 2013).

### PTBP2 post-transcriptionally controls expression of mitochondria-related genes

The discovery of irregular mitochondria in *Ptbp2* cKO spermatids prompted us to re-examine the 217 genes with altered AS mRNA isoforms in the absence of PTBP2. Notably, 24 of the 217 PTBP2-senstitive genes (11%) encode proteins that are annotated as having mitochondrial functions (Table 2), including five pairs of proteins that interact with one another (Figure 4A, green and pink lines). Differential gene expression analysis showed that the majority of these ‘mitochondria-related genes’ (79%) had no statistically significant difference in overall RNA abundance between WT and *Ptbp2* cKO testes (Table 2). This is consistent with our previous observations that most PTBP2-dependent AS regulation functions primarily to generate alternative mRNA and protein isoforms, rather than altering total transcript abundance (Licatalosi et al., 2012; Hannigan et al., 2017). To investigate whether changes in the AS RNA isoforms of mitochondrial-related genes may be primary or secondary effects of PTBP2 deletion, we asked if the corresponding pre-mRNAs were directly bound by PTBP2 in mouse testes. Using PTBP2 HITS-CLIP data from adult testes (Hannigan et al., 2017), we found that 13 of the 24 genes had biologically-reproducible PTBP2-RNA interactions within 500 nucleotides of the splice sites of either the PTBP2-dependent alternative exon and/or the flanking constitutive exons (Figure 4A, black boxes). These observations suggest that PTBP2 directly regulates the production of AS isoforms of mitochondrial-related genes in spermatogenic cells.

**Table 2.**
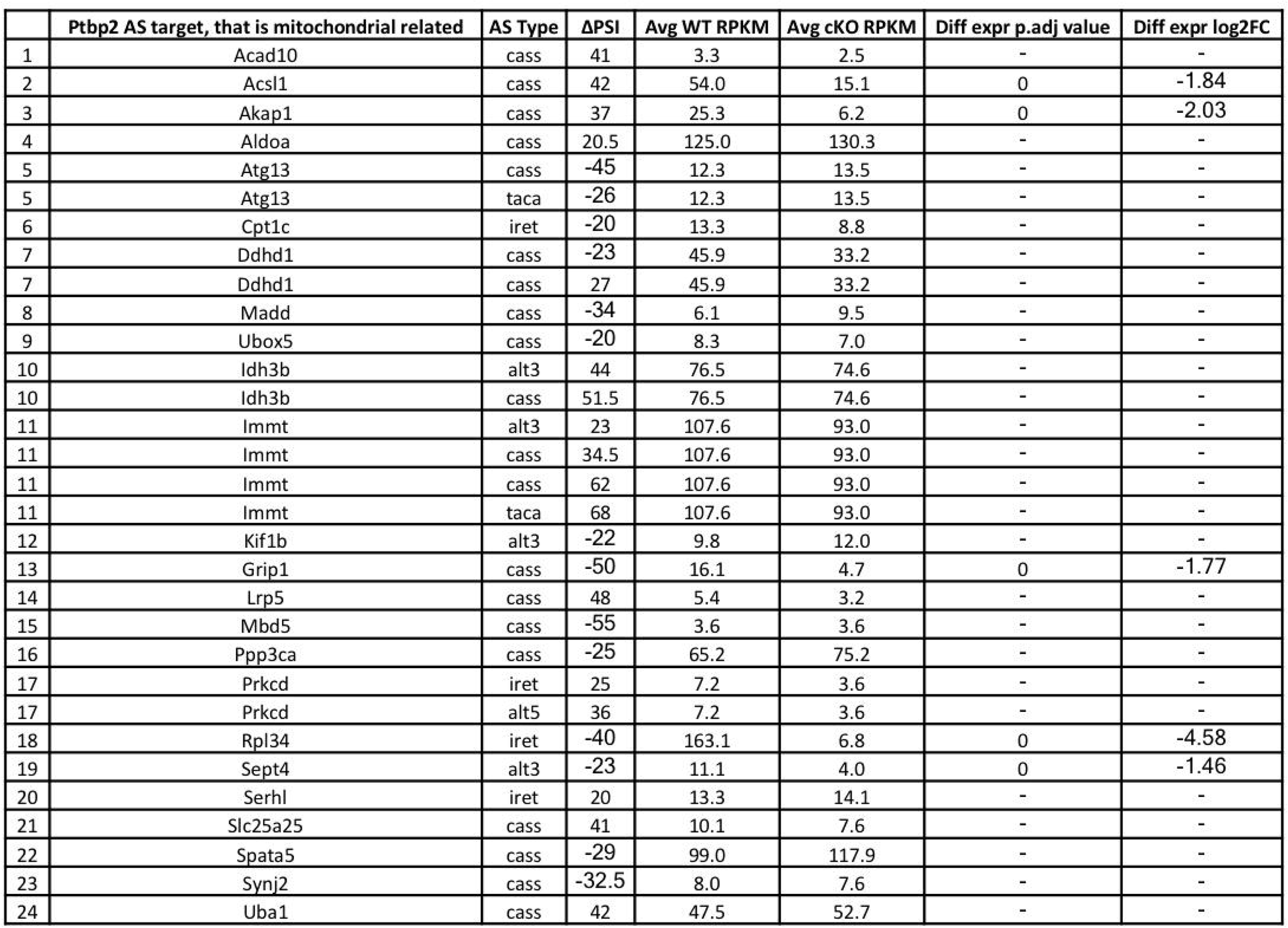
Mitochondria-related transcripts that are alternatively spliced by PTBP2. List of 24 PTBP2-sensitive transcripts that have significant AS of cassette exons (cass), tandem exons spliced in a coordinated (taca) or mutually exclusive (mutx) manner, differential 5’ and 3’ splice site selection (alt5 or alt3, respectively), and changes in intron retention (iret). ΔPSI represents that change in <u>p</u>ercent <u>s</u>pliced <u>i</u>n between WT and *Ptbp2* cKO. Gene expression data is represented as RPKM values for WT and cKO samples, and differential expression of AS transcripts (*Acsl1, Akap1, Grip1, Rpl34, Sept4*) is reported in the right two columns.

**Figure 4.**
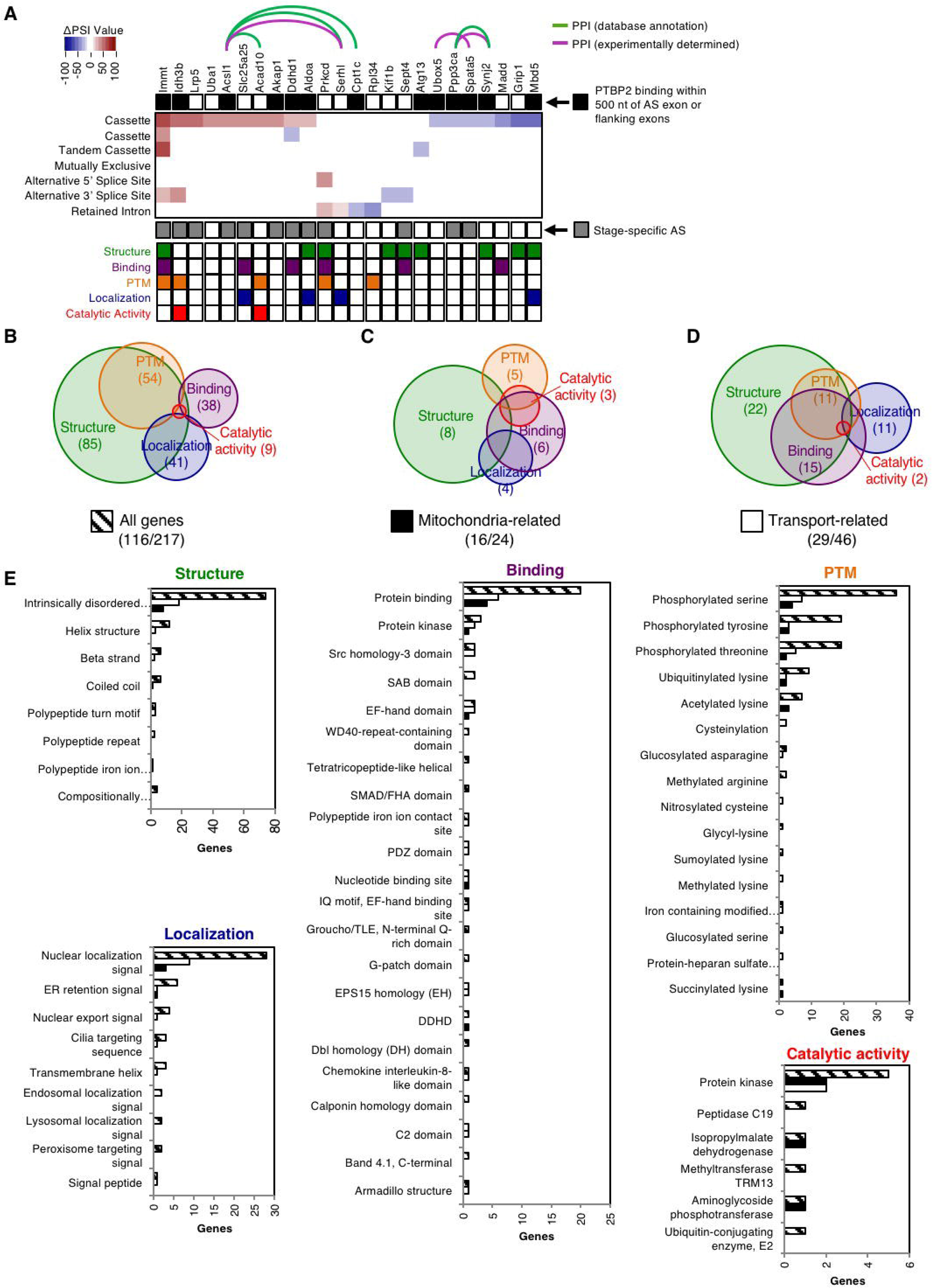
PTBP2 regulates AS isoforms of genes that control mitochondrial dynamics. A) Distribution of PSI values (change in percent spliced in between WT and Ptbp2 cKO testes) for AS events in 24 mitochondria-related genes. Red boxes represent PTBP2-enhanced splicing events, while blue boxes represent PTBP2-repressed splicing events. Genes encoding proteins involved in protein-protein interactions (PPI) are connected by green (database annotations) and/or pink (experimentally determined) lines. AS events with biologically reproducible PTBP2 binding within 500 nt of the AS exon and/or constitutive flanking exons are denoted with a black box adjacent to the affected gene. PTBP2-dependent AS events that are also regulated in a stage-specific manner are indicated by gray boxes. The bottom panel contains a summary of the Exon Ontology analysis, indicating which AS sequences impact protein structure (green), binding (purple), post-translational modification (PTM, orange), localization (blue), and catalytic activity (red). B-D) Intersection of AS genes that overlap across five main Exon Ontology functional categories for all 217 Ptbp2-sensitive genes (B), the 24 mitochondria-related AS genes (C), and genes related to transport functions (D). Panel E shows the number of genes with AS exons that encode sequences which impact specific functional features within each of the five Exon Ontology categories for all genes (hatched), mitochondria-related (black), and transport-related (white) PTBP2-sensitive genes.

A bioinformatic approach was used to investigate how PTBP2-dependent changes in AS mRNA isoforms impact the functions of the encoded proteins. Exon ontology analyses (Tranchevent et al., 2017) were performed on the coding sequences that are differentially spliced in a PTBP2-dependent manner for the 24 mitochondrial-related genes, 46 genes with roles in protein transport, and the complete set of 217 genes with PTBP2-dependent AS regulation (Supplemental File 1). Over half of the AS sequences in each gene set had annotations falling into one of five broad annotation categories (structure, localization, post-translational modification (PTM), catalytic activity, and binding), with some AS exons having annotations in multiple categories (Figure 4B-D). The most frequently observed terms associated with the PTBP2-dependent AS exons were ‘intrinsically disordered regions”, “phosphorylated serine, tyrosine, or threonine”, “protein binding”, and “nuclear localization signal” (Figure 4E). These observations indicate that PTBP2-dependent AS regulation of mRNA has wide-ranging impacts on the biochemical properties and cellular localization of the encoded proteins.

Literature searches identified multiple AS events among the 24 mitochondria-related genes which are known to alter the biochemical properties of the encoded protein. Three specific examples with proximal PTBP2-RNA interactions are highlighted here (Figure 5).

1. PTBP2-loss results in altered levels of *Ppp3ca* mRNA isoforms that contain exon 13 (Figure 5A). *Ppp3ca* encodes the phosphatase calcineurin A, which dephosphorylates DRP1 to control mitochondrial fission (Cribbs and Strack, 2007; Cereghetti et al., 2008). Splicing of *Ppp3ca* exon 13 is regulated in a tissue-specific manner and increases during postnatal development of skeletal muscle. Furthermore, functional studies in muscle indicate that AS of exon 13 impacts the intrinsic phosphatase activity of calcineurin, with reduced activity for polypeptides encoded by mRNA that contain exon 13 (Brinegar et al., 2017). Inclusion of *Ppp3ca* exon 13 is higher in spermatocytes compared to spermatogonia, and is abnormally elevated in *Ptbp2* cKO versus WT testes (Figure 5A).
2. PTBP2-loss results in altered AS isoforms of *Idh3b*, which encodes isocitrate dehydrogenase 3, a mitochondrial enzyme with a major role in the tricarboxylic acid cycle (Figure 5B). Alternative 3’ splice site selection in the last exon of *Idh3b* (exon 12) generates two IDH3B isoforms (IDH31 and IDH32) that have distinct C-terminal sequences, are expressed in a tissue-restricted manner, and have different pH optimums for IDH activity (Kim et al., 1999).
3. In the absence of PTBP2, *Immt* mRNAs have reduced splicing of exons 5 and 6. *Immt* encodes mitofilin (or Mic60), an inner mitochondria membrane protein and major component of the MICOS (mitochondrial contact site and cristae organizing system) complex (Pfanner et al., 2014) (Figure 5C). Mitofilin has roles in controlling cristae morphology and mitochondrial dynamics (John et al., 2005; Van Laar et al., 2016; Tsai et al., 2018). Encoded by *Immt* exon 6 is lysine 210 which is subject to acetylation (Choudhary et al., 2009), and 19 of 33 amino acids that comprise the first of three coiled-coil domains believed to mediate mitofilin oligomerization (Odgren et al., 1996; John et al., 2005; Tsai et al., 2018). While both exons 5 and 6 have reduced inclusion in *Immt* mRNAs in *Ptbp2* cKO testes, exon 6 shows reduced levels in spermatogonia compared to spermatocytes and spermatids, potentially reflecting different energy demands in mitotic versus post-mitotic germ cells.

**Figure 5.**
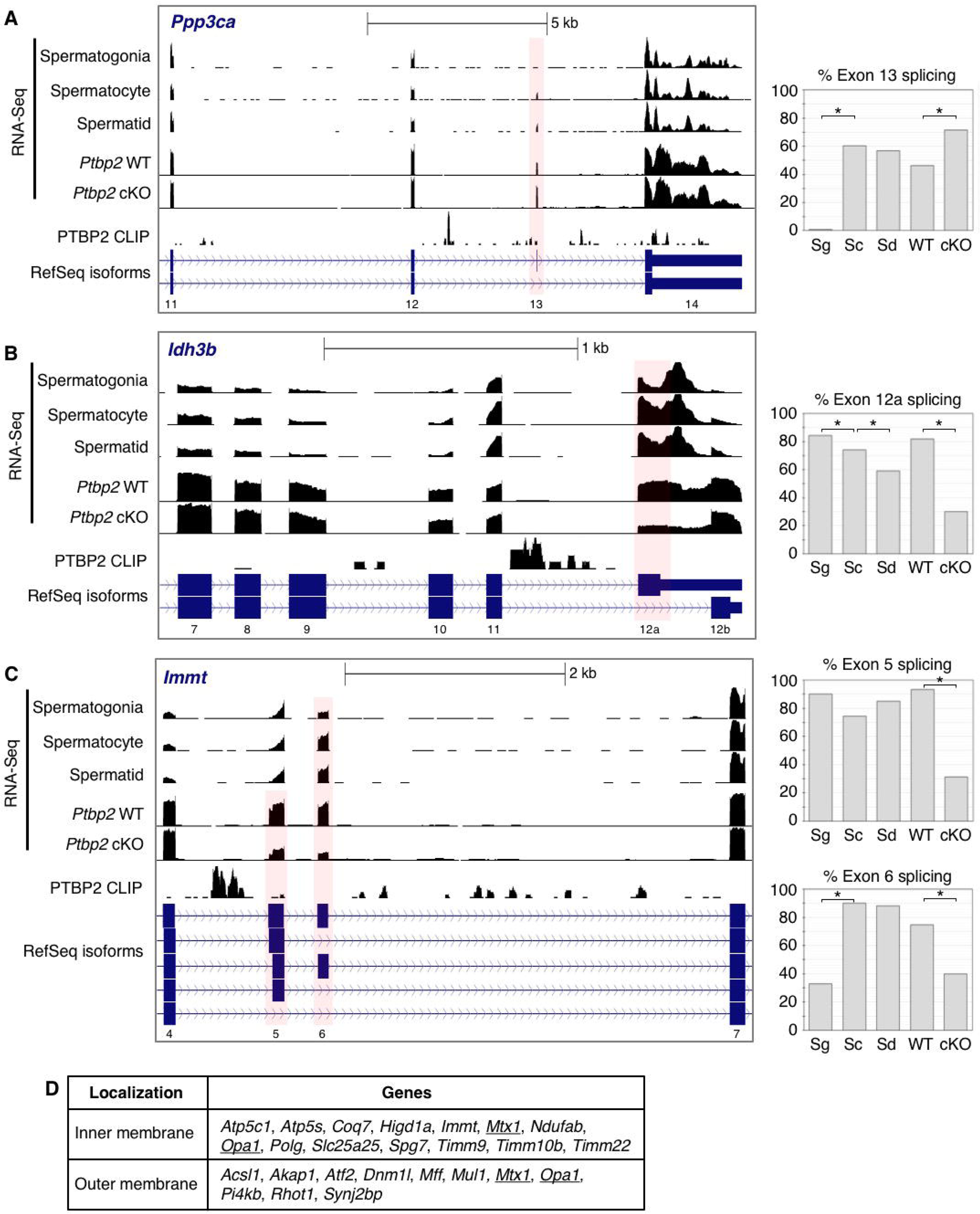
Examples of direct PTBP2 AS regulation of mitochondria-related transcripts. A-C) Three examples of mitochondria-related transcripts with PTBP2-RNA binding, and PTBP2-dependent and stage-specific regulation AS regulation. Regions of *Ppp3ca* (A), *Idh3b* (B), and *Immt* (C) genes are shown, with exons indicated below. Each panel has RNA-Seq read density data from purified spermatogonia (Sg), spermatocytes (Sc), and spermatids (Sd); RNA-Seq read density data from WT and *Ptbp2* cKO testes; and PTBP2 HITS-CLIP data. AS isoforms annotated in RefSeq are also shown. The graphs to the right show the percentage of exon inclusion for each of the indicated AS exons. Asterisks denote inclusion differences with *P*<0.002. D) List of genes with AS exons regulated in a stage-specific manner during spermatogenesis that encode proteins localized to the inner or outer mitochondrial membrane. Genes whose products localize to both membranes are underlined.

Each of these highlighted examples has an AS exon that is both PTBP2-sensitive and regulated in a stage-specific manner. Overall, in 12 of the 24 genes with PTBP2-sensitive AS events, the same exons also have differences in splicing between spermatogonia and spermatocytes, or between spermatocytes and spermatids (Figure 4A, grey boxes). Independent of PTBP2, review of our previously-defined lists of stage-specific AS changes showed that 78 of the 595 genes with splicing differences between spermatogonia and spermatocytes, and 8 of the 78 genes with splicing differences between spermatocytes and spermatids encode proteins with gene ontology annotations that include the term mitochondrion. Furthermore, “mitochondria organization” is an enriched gene ontology term for genes with AS differences between spermatogonia and spermatocytes. Among the proteins encoded by these genes are numerous proteins that localize to the inner and outer mitochondrial membrane proteins (Figure 5D).

Collectively, our observations indicate that during spermatogenesis stage-specific AS post-transcriptionally modifies expression of genes with important roles at mitochondria. How these AS isoform changes contribute, both individually and in aggregate, to the known changes in mitochondria morphology and energy metabolism in different germ cell types will require further investigation. Our results also indicate that PTBP2 is required for correct expression of AS isoforms of mRNAs that encode proteins with important roles in regulating mitochondrial dynamics and functions. Furthermore, the data suggest that an imbalance in these AS mRNA isoforms underlies the aberrant mitochondria morphology observed in *Ptbp2* cKO spermatids.

## DISCUSSION

Alternative splicing is a potent and highly regulated post-transcriptional mechanism of gene regulation that fine-tunes protein functions. Increasingly, AS is recognized as having key roles in the specialization of cell properties and as a major driver of phenotypic complexity. An emerging theme in tissue- and cell-type specific AS regulation is that particular RNA-binding proteins post-transcriptionally control sets of functionally-related genes. Well-studied examples in the nervous system include AS regulation by NOVA2 which impacts a gene network whose protein products function at neuronal synapses (Ule et al., 2005); AS regulation by RBFOX which impacts genes whose proteins localize to the axon initial segment (Jacko et al., 2018); and AS regulation by PTBP2 in the embryonic brain which is important to prevent premature neurogenesis and maintain neuronal precursors (Licatalosi et al., 2012). Similarly, regulation of AS by tissue-restricted RBPs has critical roles in development of the heart, skeletal muscle, liver, and skin (Bangru et al., 2018; Giudice et al., 2014; Bhate et al., 2015; Brinegar et al., 2017; Dittmar et al., 2012). In contrast, much less is known about the regulation and functional roles of AS in spermatogenesis, despite the long-standing recognition that unusually high levels of AS isoforms are expressed in the testis compared to other tissues (Yeo et al., 2004; Clark et al., 2007). Here we provide evidence that PTBP2-dependent AS is necessary for proper protein trafficking and mitochondrial dynamics in post-meiotic spermatids.

We hypothesized that regulation of cell polarity and protein trafficking are aberrant in *Ptbp2* cKO testes and contributes to spermatogenic arrest. This hypothesis was based on the observation that genes with altered AS mRNAs in *Ptbp2* cKO testes are disproportionately enriched for those that encode GTPases, GTPase-regulatory factors, and several proteins with roles in the selection of vesicle cargo, and vesicle formation, transport, and fusion with target membranes (Hannigan et al., 2017). Our previous observation that the Sertoli cell cytoskeleton is disorganized in *Ptbp2* cKO testes provided indirect evidence that is consistent with defects in polarity of PTBP2-deficient germ cells. In this study, we sought to more directly examine protein trafficking in *Ptbp2* cKO germ cells by TEM analysis. To do so, we took advantage of the fact that the acrosome is a large and easily identifiable spermatid-specific structure whose formation depends on continuous vesicle trafficking and fusion (Berruti, 2016). This includes endoplasmic reticulum to Golgi apparatus to acrosome transport (believed to be the main transport pathway for acrosome formation), as well as extra-Golgi trafficking via factors from endocytic and exocytotic pathways, such as AFAF and USP8 (Li et al., 2006; Berruti et al., 2010). Here, we identified several defects in acrosome formation in PTBP2-deficient spermatids. These abnormalities include attachment of multiple acrosomes to a single nucleus, impaired acrosomal membranes, and coated vesicles within the acrosome. Altogether, these observations indicate that loss of PTBP2-dependent AS regulation results in aberrant protein trafficking via transport vesicles in spermatids.

Numerous genes required for proper acrosome biogenesis have been described, including several that encode proteins involved in protein trafficking such as VPS54 (Paiardi et al., 2011), PICK1 (Xiao et al., 2009), GM130 (Han et al., 2017), GOPC (Yao et al., 2002), ATG7 (Wang et al., 2014), NHE8 (Oberheide et al., 2017), and HRB (Kang-Decker et al., 2001). The phenotypes resulting from deleting these genes are remarkably similar and generally include infertility due to the formation of spermatozoa with missshapen or round heads (globozoospermia). While acrosome formation continues in these mutants, albeit incorrectly, acrosome formation abruptly ceases during the Golgi phase in *Ptbp2* cKO spermatids. Acrosomal granules in *Ptbp2* cKO cells appear competent to attach to the nuclear membrane via the acroplaxome, and retain this activity even in multinucleated cells (evidenced by the attachment of single acrosomal granules to multiple nuclei). However, the nucleus-attached acrosomes in *Ptbp2* cKO spermatids fail to expand and flatten, suggesting a halt in protein trafficking to the acrosome. Strikingly, the few *Ptbp2* cKO spermatids that had advanced to the cap phase display overt vesicle sorting defects. Altogether, these observations suggest that mis-regulation of vesicle sorting and/or fusion underlies the failure of acrosome formation to progress beyond the Golgi phase in *Ptbp2* cKO spermatids.

Although genes required for cell polarity and protein trafficking are significantly enriched among the 217 with AS isoform imbalances in *Ptbp2* cKO testes, genes with other functions are also present. This suggests that additional cellular processes are likely dysregulated in the absence of PTBP2-dependent AS regulation. Our results indicate that mitochondrial dynamics are perturbed in *Ptbp2* cKO spermatids. This was evidenced by the accumulation of numerous elongated and dumbbell-shaped mitochondria in cKO spermatids, compared to the more spherical and evenly distributed mitochondria in WT cells. We speculated that this difference in morphology was due to an imbalance in mitochondrial dynamics (fusion and fission), which is skewed toward fission in *Ptbp2* cKO spermatids. Western blot analysis of *Ptbp2* cKO testis lysates revealed elevated levels of DRP1, a critical regulator of mitochondrial fission. While DRP1-mediated fission can be regulated by post-translational modifications (Kashatus et al., 2015; Kim et al., 2016; Perdiz et al., 2017), over-expression of DRP1 alone can result in increased mitochondrial fission (Fowler and O’Sullivan, 2016; Ma et al., 2018).

Notably, transcripts for *Dnm1l* (the gene encoding DRP1) do not appear to have differences in either steady state or AS isoform levels in *Ptbp2* cKO versus WT testes (Table 1). This suggests that PTBP2-loss indirectly results in post-transcriptional up-regulation of DRP1 that, in turn, leads to a potential increase in DRP1-oligomerization on mitochondria to promote fission. We propose that increased DRP1 results from changes in AS isoforms that encode factors that control DRP1 levels and activity. Over 10% of transcripts alternatively spliced by PTBP2 encode proteins that have mitochondrial functions, including multiple factors that control or cooperate with DRP1 (Table 2). For example, *Akap1* contains an alternative exon whose splicing is enhanced by PTBP2. AKAP1 is a member of the A-kinase anchoring proteins (AKAPS) and functions to promote localized cAMP signaling by tethering protein kinase A (PKA) to the cytosolic surface of the outer mitochondrial membrane (Flippo et al., 2018). In addition to PKA, AKAP1 can recruit other signaling molecules to mitochondria, including Src kinase and calcineurin to modulate mitochondrial function. A key substrate of AKAP1 is DRP1, where AKAP1 inhibits mitochondrial fission by promoting PKA-mediated phosphorylation of DRP1 at serine 637 (S637) (Merrill and Strack, 2014; Dagda et al., 2011; Dickey and Strack, 2011). In AKAP1-deficient mice, Flippo et al., demonstrated that deletion of AKAP1 resulted in DRP1 hyperactivation (Flippo et al., 2018). Our RNA-Seq data showed a statistically significant 2-fold reduction in *Akap1* expression in PTBP2-null mice (Table 2). Thus, the reduced splicing of alternative exon 3 and loss of *Akap1* in *Ptbp2* cKO spermatids may contribute to the observed increase in DRP1 and mitochondrial fission.

Another gene encoding a mitochondria-related protein with mis-regulated AS isoforms in *Ptbp2* cKO testis is *Ppp3ca* (Table 2). Our HITS-CLIP data (Hannigan et al., 2017) showed that PTBP2 reproducibly bound proximal to alternative exon 13 to repress its splicing (Figure 5A). Notably, *Ppp3ca* encodes the alpha isoform of the calcineurin catalytic subunit. Calcineurin is a ubiquitously expressed protein phosphatase that is dependent on calcium and calmodulin binding in order to function as a signaling regulator for mitochondrial dynamics (Pfluger et al., 2015); and as mentioned above, calcineurin is also a target of AKAP1. Calcineurin promotes mitochondrial fission by dephosphorylating the PKA-dependent site S637. Previous studies have shown that calcineurin physically interacts with DRP1 and that inhibition of this phosphatase blocks DRP1-dependent fission of mitochondria (Cereghetti et al., 2008; Pennanen et al., 2014). DRP1-mediated fission can also be influenced by protein kinase C delta (PRKCD) – whose transcript is also alternatively spliced by PTBP2 (Table 2) and interacts with PPP3CA (Figure 3). Like AKAP1 and calcineurin, PRKCD physically interacts with DRP1 (Zaja et al., 2014), yet mediates an increase in mitochondrial fragmentation by phosphorylating DRP1 at serine 579 (S579) (Qi et al., 2011). Importantly, previous studies showed that inhibition of PRKCD reduced mitochondrial fragmentation, fission, and DRP1 levels (Qi et al., 2011). Thus, it is possible that the alternative splicing of *Prkcd* mRNA by PTBP2 may change the activity of this kinase, impacting its ability to phosphorylate DRP1 and activate mitochondrial fission.

Additional examples highlighting how mis-regulation of AS isoforms can impact mitochondria include PTBP2-dependent splicing differences in *Synj2* and *Spata5* mRNAs. SYNJ2 (Synaptojanin-2) is a mitochondrial-shaping protein involved in mitochondrial aggregation, membrane trafficking and signaling transduction (Rocha et al., 2017). SPATA5 was originally described as a spermatogenesis-associated factor with two-conserved ATPase modules and a mitochondria localization signal (Liu et al., 2000). In neurons, SPATA5 had a critical role in mitochondrial morphology, with *Spata5* deficiency resulting in decreased mitochondrial fusion (Puusepp et al., 2018). Collectively, our observations indicate that PTBP2-dependent AS regulation has an important role in maintaining proper mitochondrial dynamics in post-meiotic spermatids.

Assessing how changes in individual mRNA isoforms contribute to phenotype is a major challenge in the study of AS and its functional roles *in vivo*. While some cellular processes are likely to be more sensitive to certain AS isoform changes, a recurring theme in cell- and tissue-specific AS regulation is that networks of co-regulated AS isoforms work in combinatorial manner to shape cellular properties and expand phenotypic complexity (Licatalosi and Darnell, 2010; Baralle and Giudice, 2017). Our observation that several genes encoding mitochondrial proteins are co-regulated by stage-specific AS suggests that post-transcriptional regulation may contribute to changes in mitochondria morphology and energy metabolism during spermatogenesis (Meinhardt et al., 1999).

Interestingly, intrinsically disordered regions are particularly impacted by PTBP2-dependent AS regulation in spermatogenic cells. Intrinsically disordered proteins (IDP) play an important role in cellular signaling and are subject to post-translational modifications given their enhanced flexibility and conformational plasticity. While several different enzymes can modify IDPs, phosphorylation sites are located predominately in intrinsically unstructured regions (Wright and Dyson, 2015), likely supporting the enrichment of tyrosine and serine phosphorylation functional features among our list of PTBP2-sensitive transcripts. Intrinsically disordered regions introduced by alternative splicing are also known to increase protein-protein interactions by serving as a scaffold for multiple protein partners (Babu, 2016), thereby contributing to cellular complexity.

In summary, our study demonstrates that germ cell-specific loss of PTBP2, a tissue-restricted AS regulator, leads to defects in protein transport and mitochondrial defects consistent with increased fission. Incorporating observations from ultrastructural analyses and transcriptome-profiling data indicates that PTBP2 has important roles in maintaining the correct levels of AS mRNA isoforms necessary for proper protein trafficking and regulation of mitochondria in spermatogenesis. Collectively, our findings illustrate the importance of AS regulation and PTBP2 in mammalian germ cell development.

## ACKNOWLEDGEMENTS

We are grateful to Alan Tartakoff for reviewing electron micrographs.

---

**Supplemental File 1.** Excel file with results from Exon Ontology analysis of the 217 genes with PTBP2-dependent AS regulation. Each sheet corresponds to one of the five annotation categories (structure, binding, post-translational modification (PTM), localization, and catalytic activity). Within these sheets are subsets of the 217 AS genes that were assigned Exon Ontology features, and whether or not the AS gene encodes a protein with function(s) related to mitochondria and/or transport.

## MATERIALS AND METHODS

### Animal and Tissue Collection

WT and *Ptbp2* cKO C57BL/6 littermates were generated by crossing *Stra8-iCre^+/+^; Ptbp2^ΔE4/+^* males with *Ptbp2^flox/+^* females, as previously described (Zagore, 2015). For all procedures, mice were anesthetized using isoflurane inhalation and sacrificed by decapitation. Testes were collected and decapsulated in Gey’s Balanced Salt Solution on ice. Institutional Animal Care and Use Committee at Case Western Reserve University (CWRU) approved all animal protocols.

### Bright Field and Electron Microscopy

Small pieces of tissue collected from P25 and P30 testes (WT and *Ptbp2* cKO in replicate) were fixed by immersion in triple aldehyde-DMSO (Kalt and Tandler, 1971). After rinsing in distilled water, samples were postfixed in ferrocyanide-reduced osmium tetroxide. Another water rinse was followed by an overnight soak in acidified uranyl acetate. After rinsing in distilled water, the tissue blocks were dehydrated in ascending concentrations of ethanol, passed through propylene oxide, and embedded in Poly/Bed resin (Fujioka et al., 2012). Thin sections were sequentially stained with acidified uranyl acetate followed by a modification of Sato’s triple lead stain (Hanaichi et al., 1986) and examined in an FEI Tecnai T12 electron microscope equipped with a Gatan single tilt holder and a Gatan US4000 4kx4k CCD camera.

### Immunoblotting

WT and *Ptbp2* cKO testes from P24 (in replicate) and P30 mice were lysed in buffer containing 150 mM NaCl, 20 mM Tris-HCl pH 8.0, 1% TritonX-100, 0.1% SDS, and a protease inhibitor tablet (Roche 10481700). Proteins were transferred onto polyvinylidene fluoride (Millipore) membranes and probed with the following antibodies: α-DRP1 (sc-101270, 1:1000); α-PTBP2 (ab154787, 1:5000); α-HSP90 (BD 610418; 1:1000). Isotype-matched, horseradish peroxidase-conjugated secondary antibodies (Invitrogen) were used, followed by chemiluminescence detection.

### High-throughput data analyses

RNA-Seq of WT and *Ptbp2* cKO testes, as well as PTBP2 HITS-CLIP data of WT mice were previously described (Hannigan et al., 2017). Protein features encoded by Ptbp2-sensitive alternative exons were identified by the publically available computational resource, “Exon Ontology” (Tranchevent et al., 2017). For this analysis, we converted mm9 coordinates corresponding to the 257 AS events changed in *Ptbp2* cKO testes to hg19 annotations. Of the 257 events, 244 coordinates were successfully converted to the human genome. Protein-protein interaction analysis of the 24 Ptbp2-sensitive AS targets with mitochondria-related functions was performed using the STRING database (Szklarczyk et al., 2017).

